# Habitat-Net: Segmentation of habitat images using deep learning

**DOI:** 10.1101/483222

**Authors:** Jesse F. Abrams, Anand Vashishtha, Seth T. Wong, An Nguyen, Azlan Mohamed, Sebastian Wieser, Arjan Kuijper, Andreas Wilting, Anirban Mukhopadhyay

## Abstract

Understanding environmental factors that influence forest health, as well as the occurrence and abundance of wildlife, is a central topic in forestry and ecology. However, the manual processing of field habitat data is time-consuming and months are often needed to progress from data collection to data interpretation. Computer-assisted tools, such as deep-learning applications can significantly shortening the time to process the data while maintaining a high level of accuracy. Here, we propose Habitat-Net: a novel method based on Convolutional Neural Networks (CNN) to segment habitat images of tropical rainforests. Habitat-Net takes color images as input and after multiple layers of convolution and deconvolution, produces a binary segmentation of the input image. We worked on two different types of habitat datasets that are widely used in ecological studies to characterize the forest conditions: canopy closure and understory vegetation. We trained the model with 800 canopy images and 700 understory images separately and then used 149 canopy and 172 understory images to test the performance of Habitat-Net. We compared the performance of Habitat-Net with a simple threshold based method, a manual processing by a second researcher and a CNN approach called U-Net upon which Habitat-Net is based. Habitat-Net, U-Net and simple thresholding reduced total processing time to milliseconds per image, compared to 45 seconds per image for manual processing. However, the higher mean Dice coefficient of Habitat-Net (0.94 for canopy and 0.95 for understory) indicates that accuracy of Habitat-Net is higher than that of both the simple thresholding (0.64, 0.83) and U-Net (0.89, 0.94). Habitat-Net will be of great relevance for ecologists and foresters, who need to monitor changes in their forest structures. The automated workflow not only reduces the time, it also standardizes the analytical pipeline and, thus, reduces the degree of uncertainty that would be introduced by manual processing of images by different people (either over time or between study sites). Furthermore, it provides the opportunity to collect and process more images from the field, which might increase the accuracy of the method. Although datasets from other habitats might need an annotated dataset to first train the model, the overall time required to process habitat photos will be reduced, particularly for large projects.

## 1. INTRODUCTION

Understanding of the intricacies of natural forest ecosystems is important to better manage and protect them. In both ecology and forestry there is a huge need for high quality information about the forest structure to help understand the spatio-temporal changes of forest habitat (Stojanova *et al.*, 2010). Detailed, large-scale knowledge about forest habitat and structure and how forests respond to anthropogenic disturbance will improve our ability to mitigate the effects of disturbances. Canopy closure and understory vegetation density are measurements that are commonly used in forest and land use research, monitoring, management and planning (Jennings, Brown, & Sheil, 1999).

Canopy closure is the proportion of sky hemisphere obscured by vegetation when viewed from a single point (Jennings, Brown, & Sheil, 1999). More rigorously defined, canopy closure is defined as the percent forest area occupied by the vertical projection of tree crowns (Paletto & Tosi, 2009). The canopy closure metric is an important part of forest inventories (Korhonen *et al.*, 2006; Chopping *et al.*, 2008), linked with canopy architecture, light regimes, solar radiation and leaf area index estimates in forest ecosystems. It is useful for wildlife habitat assessment and monitoring (Paletto & Tosi, 2009) and is often used as a multipurpose ecological indicator (Korhonen *et al.*, 2006). For forestry practitioners, measurements of the forest canopy serve as one of the chief indicators of the microhabitat within the forest (Jennings, Brown, & Sheil, 1999). The forest canopy affects plant growth and survival, hence determining the nature of the vegetation, and wildlife habitat. Characterization of the understory vegetation is of equal importance in forestry as it plays a central role in forest ecosystem structure and composition (Russell et al., 2014). Understory vegetation provides key elements (and indicators) for biodiversity, nutrient cycling and forest fuel loads, and shape overstory tree structure and diversity (Halpern and Spies, 1995, Legare et al., 2002; Gilliam, 2007; Russell et al., 2014). Understory vegetation has become a fundamental component of forest site classifications (Bergès Gégout, & Franc, 2006) and the status of the understory composition and structure is a critical indicator of the condition of the forest (D’Amato, Orwig, Foster, 2009).

For ecologists and wildlife managers there is also a great need to understand factors influencing the occurrences and habitat preferences of species (Cristescu & Noyce, 2013), as the ecology of many species is poorly understood hindering their effective management. Therefore, accurate and quick habitat characterizations of the surveyed sites are needed (Zeng *et al.*, 2013). Canopy closure and understory vegetation are among the factors that are known to influence species occurrence at a site (Vickers & Palmer, 2000; Brenes-Arguedas *et al.*, 2011).

Despite the importance of quantitative estimates of canopy closure and understory vegetation density, there are no efficient ways to obtain these values. A plethora of different techniques have been developed to quantify forest canopy (Jennings, Brown, & Sheil, 1999). Conventionally, information on canopy closure was collected using a spherical densiometer (Jennings, Brown, & Sheil, 1999). This technique is very labour intensive and requires researchers to spend a lot of time in the field. Currently, canopy closure information is often obtained through manual processing of color digital canopy photographs to binary images of vegetation and sky or through simple thresholding methods (Jonckheere *et al.*, 2005; Nobis, & Hunziker, 2005), which is less time consuming in the field but requires lots of manual processing at the computer. One of the traditional methods to characterize understory vegetation in the field is through the use of cover boards. Here an observer visually estimates the relative proportion of a board of known dimensions that is being obscured by vegetation from a given vantage point (Jones, 1968; Nudds, 1977). The subjectivity inherent to the visual estimation by an observer is a widely acknowledged limitation of both the spherical densiometer and cover board field methods (Limb *et al.*, 2007; Morrison, 2016). Recently, the workflow for the processing of both canopy and understory images has become digital due to advancements in digital photography and image processing of digital vegetation photographs (Marsden et al, 2002; Jorgensen et al., 2013).

Currently, due to logistical and analytical challenges, and time consuming manual processing of field data, several months, and sometimes up to years, are needed to progress through the stages of data collection, data processing, and interpretation. The processing of the habitat photograph datasets, which are usually large, is presently done using manual methods or simple thresholding methods. This is highly unsatisfactory and prevents timely evidence-based, effective action in both forestry and conservation applications. Fast, reliable, and automated computer-assisted tools are therefore needed to describe the habitat immediately after data collection. Using tools such as advanced machine learning, which have gained popularity when solving many data driven tasks in other fields, is one option to overcome the time demand associated with habitat interpretation. Some more advanced techniques for the processing of habitat images, some of which exploit machine learning, do exist, including traditional LiDAR-based and 3D image based techniques to segment canopy and understory images (Stojanova *et al.*, 2010; Tao *et al.*, 2015; Hamraz *et al.*, 2017). These methods however, require expensive equipment to collect LiDAR or 3D images and seldomly are these data available for forest inventories or ecological studies. Therefore most projects still rely on either labour intensive field methods, such as the use of spherical densiometers, or on digital habitat photos, which later need manual processing. Thus, the processing of thousands of simple digital habitat photographs desperately requires advances in automated workflows.

Recently, computer vision has made an inroads to the ecological domain. There have been limited attempts to use machine learning methods for automated interpretation of forest habitat images, such as canopy closure photographs (Levner & Bulitko, 2004; Zhao *et al.*, 2010; Erfanifard, Khodaei, & Shamsi, 2014; Ahmed *et al.*, 2015). Compared to these early studies, more advanced deep learning techniques have been developed, particularly in the field of medical imaging research. These methods have proved to be superior to earlier techniques (Litjeans *et al.*, 2017), significantly shortening the time and increasing the accuracy of the data interpretation and data processing. Specifically, Convolutional Neural Networks (CNNs), which are deep feedforward neural networks that are inspired by the visual cortex of the human eye and allow computers to ‘see’, have been shown to outperform many state-of-the-art methods in various visual computing tasks across different domains (Krizhevsky, Sutskever, & Hinton, 2012). CNNs are used to recognize images by processing the original image through multiple layers of feature-detecting “neurons”. Each layer is designed to detect a specific set of features such as lines, or edges. Increasing the number of layers (typical CNNs use anywhere from 5 to 25 layers or more) allows the CNN to detect more complex features enabling it to recognize the object in the image. Biomedical images generated by a wide range of medical procedures, such as MRI and CT scans, have complex textural patterns and are limited to small annotated datasets making it difficult to apply machine learning techniques and classic CNNs. However, the U-Net model helped overcome these challenges in biomedical image segmentation (Ronneberger, Fischer, & Brox, 2015). This is due to the network architecture that consists of a contracting path, in which the spatial information is reduced while feature information is increased, and an expansive path, which combines the feature and spatial information from the contracting path.

In practice, the application of CNNs to medical imaging is very similar to their application in the field of ecology. Similar to medical images, canopy and understory images contain a lot of color and textural information. In this work we focus on using computer vision methods to automatically extract the relevant information about canopy closure and understory vegetation density from in-situ digital photographs. We propose Habitat-Net, a novel deep learning method based on U-Net, to segment in-situ habitat images of forests. We extend the U-Net architecture to a new domain and improve on the performance of U-Net by implementing Batch Normalization. Our proposed framework has been designed to feed color forest habitat (both canopy and understory) images as input to the network and, after multiple convolutions, generates a binary segmentation raster of the original image. The entire pipeline of the proposed method has been designed to work automatically without any user interaction. Our only assumption is that during model training the input images are accompanied by respective annotated images.

## 2. METHODS

### 2.1. Dataset Description

We conducted standardized vegetation surveys in Sabah, Malaysian Borneo between 2014 and 2016. We collected a total of 949 canopy (128 x 128 pixels) and 872 understory vegetation (256 x 160 pixels) photographs that are used in this study. All photos were taken using the built-in camera in GPS unit (Garmin® model 62sc). To collect our canopy dataset in the field, we established a 20 x 20 m grid around the center point of our survey station, which was located halfway between the two camera traps. The grid was positioned along the north-south, east-west axes. We took canopy photographs at the centerpoint and the NW, NE, SW, and SE corners of the plot. All canopy photos were taken at an angle of approximately 90 degrees (directly overhead). The understory dataset was collected by taking photos of a 1.5 x 1.0 m orange fly-sheet positioned 10 m in each cardinal direction while standing at the centerpoint of the survey grid. The vegetation covering the flysheet is used to estimate the understory vegetation density. The orange sheet used during data collection separates the understory areas from the background providing a means to segment the understory images. The photographs range in complexity from a completely uncovered orange sheet with no understory visible (the reference is an entirely white image) to images where the orange sheet is completely covered due to dense understory structure (reference segmentation image is an entirely black image).

### 2.2. Manual segmentation Deriving canopy closure and vegetation density from field photographs

We used the free and open source image manipulation software Gimp to process the canopy and understory images. We set color thresholds and used the binary indexing feature in Gimp to convert the color canopy images into binary black and white images with black representing foliage and white sky (Fig. 1). The processing of the understory vegetation photos followed the same basic workflow, segmenting a binary image, but included an additional preprocessing step in which we cropped the image to the extent of the orange flysheet that was being photographed from a 10 m distance. Similar to the canopy closure this workflow resulted in a binary (black and white) raster with black representing vegetation and white the orange flysheet (gaps in understory vegetation). Canopy closure and vegetation density was then calculated from the classified binary images by automatically counting black (vegetation) and white (non-vegetation) pixels using the following R script:

**Figure 1:**
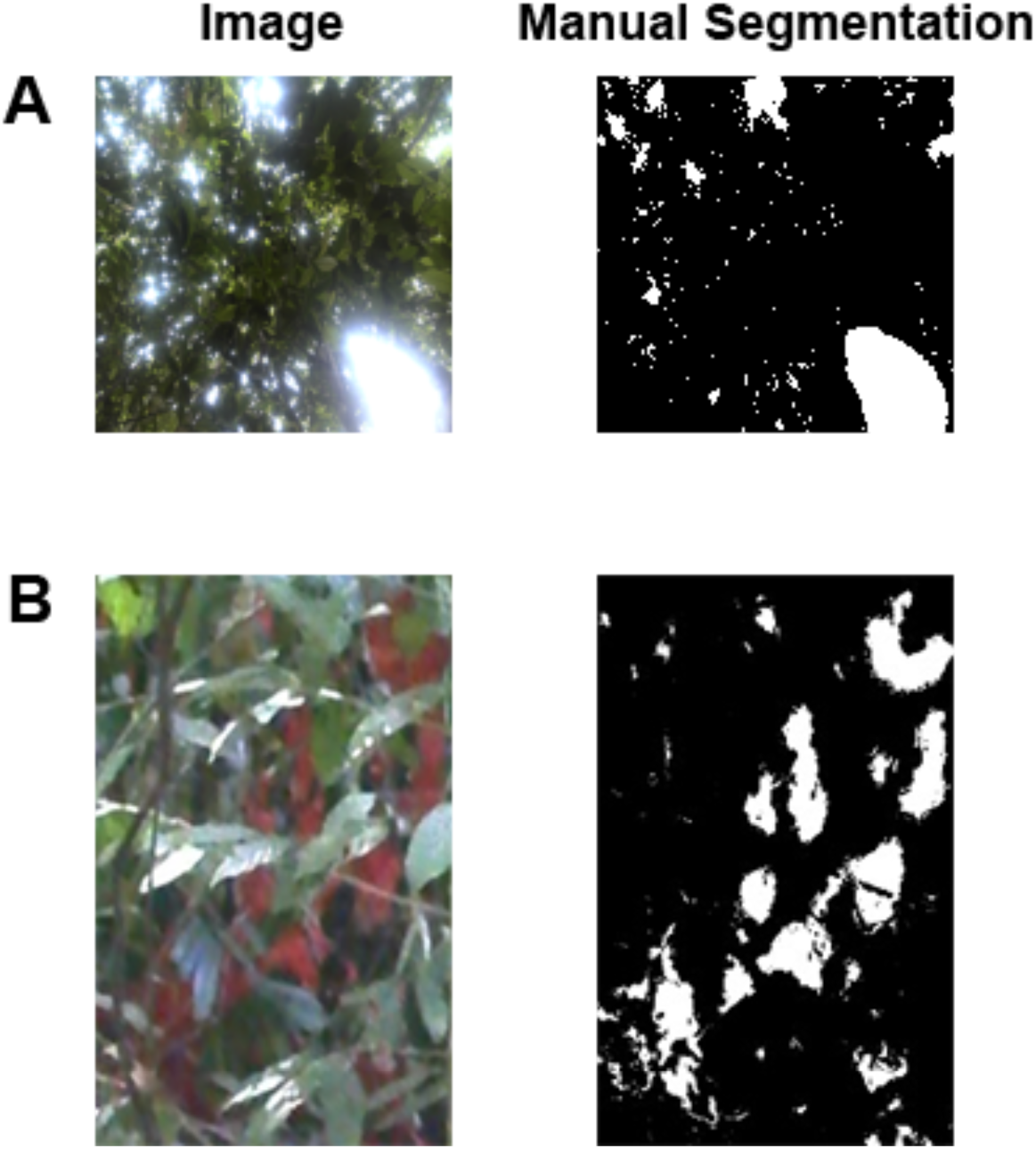
The first column contains the digital color images from photographs of (A) canopy and (B) understory taken in the field. The second column shows the manually segmented binary images of the same (A) canopy and (B) understory photographs.

~~~
*# load image as raster and convert to matrix*
r <-raster(files.tmp[j])
r.matr <-as.matrix(r)
*# set threshold value to split vegetation and sky/empty space: range = 0 - 256. 128 is medium grey (needed because jpg creation from binary image introduces some artefacts which are then made binary again)* thresholdValue = 128,
*# consider as vegetation all pixels with value < thresholdValue*
fraction_vegetation <-as.numeric(mean(r.matr < thresholdValue))
*# the rest is sky/empty space*
fraction_sky <-1 fraction_vegetation
~~~

#### Second manual segmentation

In order to compare the performance and consistency of manual segmentation by different researchers, a second independent researcher performed the manual segmentation on the test set of canopy and understory images.

### 2.3. Simple thresholding

Image thresholding is a simple, yet effective, image segmentation technique that partitions an image into a foreground and background. We ran a simple thresholding algorithm (scripts available in the Supplementary Material) to convert our color images to monochrome images. We first converted field images to grayscale and then used Otsu’s method to automatically reduce the grayscale image to a binary image (Otsu, 1979). In short, in Otsu’s method we assume that the image contains two classes of pixels following a bimodal histogram (foreground pixels and background pixels). We then calculate the threshold that minimizes the intra-class variance (the variance within the class), defined as a weighted sum of variances of the two classes by iterating through all possible threshold values. We implemented this in Python version 3.7.1 using the *threshold_otsu* function from the package *scikit_image* version 0.14.1 (van der Walt *et al.*, 2014).

### 2.4. Habitat-Net

The Habitat-Net architecture (Fig. 2) is based on the U-Net (Ronneberger, Fischer, & Brox, 2015) convolutional network which provides pixel level localization by combining high resolution features with upsampling layer outputs. The U-Net model architecture has a large number of feature channels that allow the network to propagate context information to higher resolution layers. As a consequence, the expansive path is symmetric to the contracting path and yields a U-shaped architecture (Ronneberger, Fischer, & Brox, 2015).

**Figure 2:**
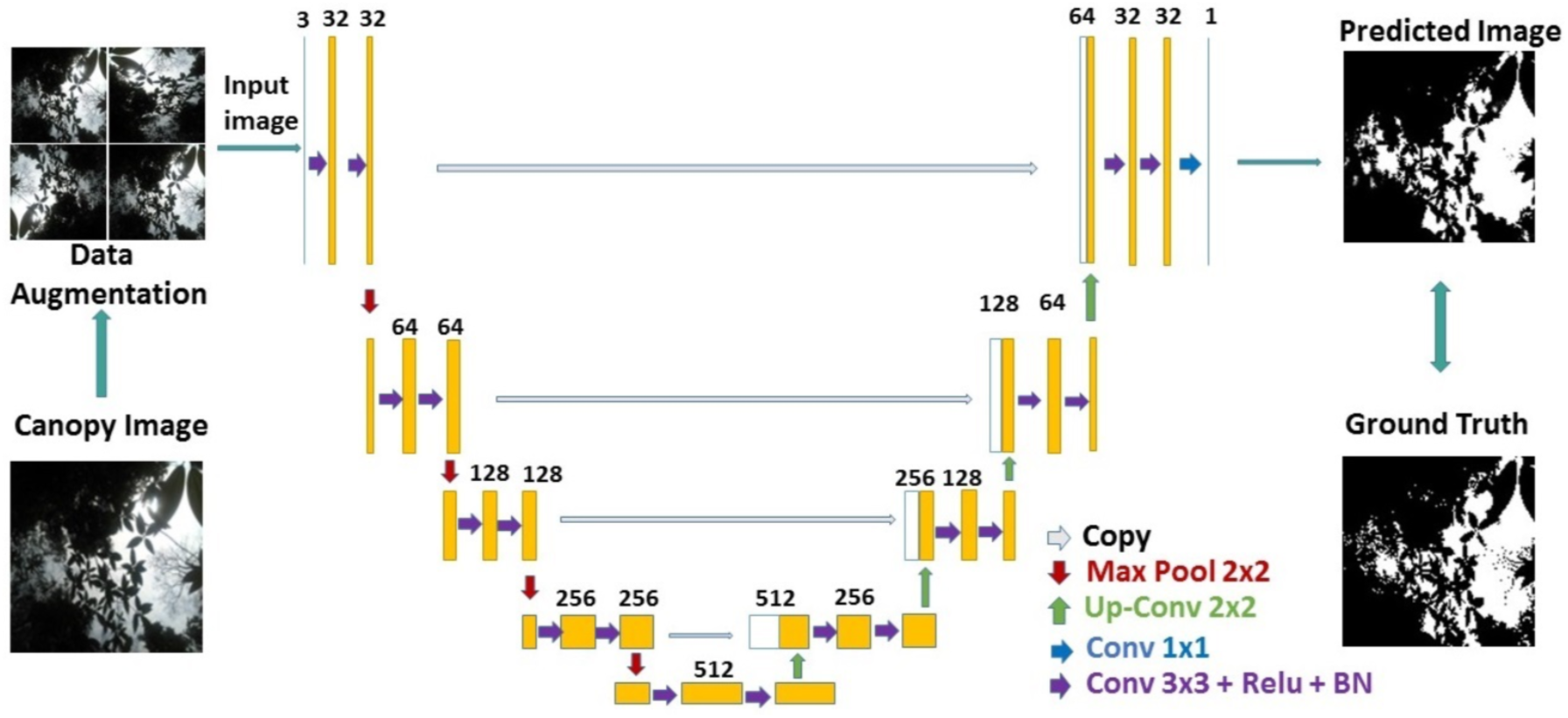
Habitat-Net architecture based on U-Net (Ronneberger, Fischer, & Brox, 2015).

The Habitat-Net consists of multiple 3 × 3 convolutions followed by a non-linear activation using rectified linear unit (ReLU). The use of a small filter size helps to capture finer details of the image. To achieve a numerically stable training procedure, we incorporate a batch normalization (BN) layer (Ioffe & Szegedy, 2015) after every convolution layer as a novel design choice in Habitat-Net. The batch normalization layer reduces the internal covariate shift, which can boost segmentation performance and helps to make training more resilient to the parameter scale using mini-batches. Batch normalization further reduces overfitting to the minimum extent and replaces the need for a dropout layer in most cases (Srivastava *et al.*, 2019). Every batch normalization layer is followed by a 2 × 2 max pooling operation, which operates independently on every depth slice of the input matrix and resizes it spatially using the max operation. We increased feature channels by an order of two at every downsampling step and halved the feature channels at each upsampling step. We used a zero-padding hyperparameter to control the spatial size of the output volumes (source code is provided in: https://github.com/Kanvas89/Habitat-Net).

Our final network includes a total of 33 convolutions including the final 1 × 1 out convolution layer. The presence of many convolution layers helps to achieve better model accuracy (Szegedy *et al.*, 2015). The only trade-off of this increased accuracy is that the network requires more time and resources to converge. We use a stochastic gradient descent (SGD) optimizer with time-based decay, which handles limited datasets well. SGD performs better in vision-based machine learning tasks and generalizes quicker than other adaptive methods such as Adam, RMSProp, and AdaGrad (Wilson *et al.*, 2017). SGD with momentum helps the parameter vector to build up velocity in any direction with a constant gradient descent so as to prevent oscillations, which leads to faster-convergence (Qian, 1999; Sutskever *et al.*, 2013). The final output layer returns a one-channel grayscale prediction image of the input habitat (canopy and understory) image. The binary raster is then run through the same R function that is used for the manual processing. All the experiments have been run using Keras 2.2.2 with TensorFlow 1.10.0 using python 3.5 on a system with a Nvidia 1080 Ti GPU.

#### 2.4.1 Network training and testing

Here, we test our method using the canopy closure and understory density datasets. The training datasets consist of a pair of images (either canopy or understory), the image to be segmented and a manual segmentation raster drawn by an expert to be used as a reference to train the model (Fig. 1). Deep neural networks typically perform better with more training data. Models trained on small datasets do not generalize well and suffer from overfitting (Perez & Wang, 2017). When a limited number of images with complex textural and color patterns are available to train the model, it is imperative to exploit data augmentation to increase the total number of training images. However, the non-linear transformations used for augmenting cell images in U-Net (Ronneberger, Fischer, & Brox, 2015) can not be incorporated directly in Habitat-net due to domain specific properties. We address this issue by artificially inflating the number of training images through rotations and reflection of the image (Supplementary Figs. S1 & S2).

Of 949 color canopy images, the training consists of 800 image pairs (image and respective reference segmentation raster). After applying the data augmentation technique our training dataset has a total of 4000 image pairs. We use 3400 images for the training dataset and the remaining 600 for validation during training. Of the 872 understory vegetation photos 700 images (after augmentation 3500 images) were used in the training dataset. Similar to the manual processing, it was necessary to perform the preprocessing step of cropping the understory images to the extent of the orange flysheet as this could not be automated. The code and the trained weights for Habitat-Net can be accessed at https://github.com/Kanvas89/Habitat-Net. As the network converged faster and variance was lower when batch normalization was used after each convolution layer (Supplementary Fig. S3) we applied batch normalization in all results presented below.

After training Habitat-Net, we tested its performance on the remaining 149 canopy images and 172 understory images (15% of the total dataset). To evaluate the performance of Habitat-Net we used overlap ratio measures (Jaccard 1907; Dice 1945; Sørensen, 1948), which quantify the degree of similarity between two objects. In our case they are an indicator of the overlap between our manually segmented reference images and those generated by the Habitat-Net model. We report both the Dice coefficient and Jaccard index as segmentation quality metrics to evaluate the performance of Habitat-Net. This is because although the Dice coefficient and Jaccard index are similar, the Jaccard index is numerically more sensitive to mismatch when there is reasonably strong overlap. As the Jaccard index penalizes single instances of bad classification more than the Dice coefficient, results of the Dice coefficient typically “look nicer” because they are higher for the same pair of segmentations and thus the Dice coefficient index is currently more popular than the Jaccard index.

## 3. RESULTS

Based on researcher experience, the manual processing of a canopy closure image requires about 45 seconds per image, while the processing of an understory image requires about 65 seconds per image (this largely varies depending on the level of experience of the individual researcher and quality of the images). Habitat-Net reduces total processing time to around 15 milliseconds per image for both canopy and understory images. Similar simple thresholding significantly reduces the time for 1 image to less than 1 second. For a typical dataset of around 400 images, both Habitat-Net and simple thresholding reduce total processing time to seconds compared to the manual processing for which 5 hours (canopy images) or even 7.5 hours (understory images) were needed (Tables 1 and 2). Visual inspection of the segmentation results (Fig. 3) from the three automated methods indicate that Habitat-Net and U-Net (both machine learning methods) outperform the simple thresholding method for the canopy images. However, the improved performance for the understory images is not as obvious in many cases. However the quantitative assessment of the performance of the different methods using the Dice and Jaccard similarity scores reveal the greatest accuracy of Habitat-Net (Fig. 4, Tables 1 & 2). Particular for the canopy images the similarity scores of the Habitat-Net were with 0.94 Dice and 0.88 (Jaccard) much higher than for the other methods, including the U-Net upon which Habitat-Net is based (Table 1). The differences for the understory images between the different methods were less strong, as all methods had higher similarity scores. However again Habitat.-Net outperformed the other methods with a Dice score of 0.95 and a Jaccard index of 0.92 (Table 2). Although Habitat-Net generally outperformed other segmentation methods, there were a few extreme outliers produced. We visually inspected the color photographs of the images with outlying similarity scores for Habitat-Net (Fig. 4). For canopy images, the images with the poorest similarity scores are “speckled” images that contain many small (sometimes single pixel) openings in the canopy vegetation. However, for the canopy images the minimal similarity score is 0.75, indicating that even when Habitat-Net performs poorly, the resulting segmentation is still acceptable. There are, however, some more extreme outliers present for the understory images. These images are very dark, blurry, and either all or almost all of the orange flysheet is covered, leaving only small openings in the vegetation. Although we inspected the problems in images which were outliers in the Habitat-Net analysis similar outliers, very likely in the same images were found in all methods, even in the manual processing of a second researcher.

**Table 1:** Mean processing time per photo and similarity scores for the canopy dataset using four methods.

| Method | Processing time (seconds) | Dice coefficient |  |  | Jaccard index |  |  |
| --- | --- | --- | --- | --- | --- | --- | --- |
|  |  | Mean | SD | Median | Mean | SD | Median |
| Manual | 45 | 0.84 | 0.16 | 0.86 | 0.74 | 0.19 | 0.76 |
| Simple thresholding | 0.103 | 0.64 | 0.17 | 0.61 | 0.49 | 0.19 | 0.45 |
| U-Net | 0.015 | 0.89 | 0.07 | 0.89 | 0.81 | 0.11 | 0.80 |
| Habitat-Net | 0.015 | 0.94 | 0.06 | 0.95 | 0.88 | 0.09 | 0.90 |

**Table 2:**
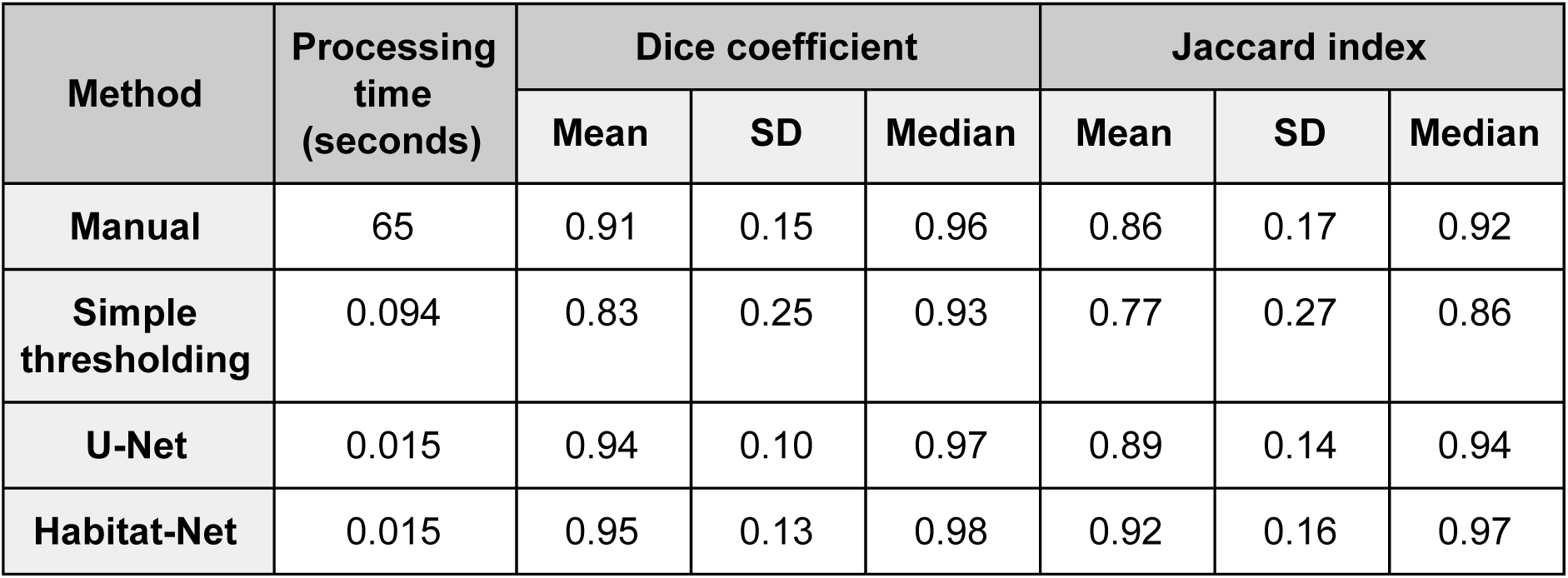
Mean processing time per photo and similarity scores for the understory vegetation dataset using four methods.

| Method | Processing time (seconds) | Dice coefficient |  |  | Jaccard index |  |  |
| --- | --- | --- | --- | --- | --- | --- | --- |
|  |  | Mean | SD | Median | Mean | SD | Median |
| Manual | 65 | 0.91 | 0.15 | 0.96 | 0.86 | 0.17 | 0.92 |
| Simple thresholding | 0.094 | 0.83 | 0.25 | 0.93 | 0.77 | 0.27 | 0.86 |
| U-Net | 0.015 | 0.94 | 0.10 | 0.97 | 0.89 | 0.14 | 0.94 |
| Habitat-Net | 0.015 | 0.95 | 0.13 | 0.98 | 0.92 | 0.16 | 0.97 |

**Figure 3:**
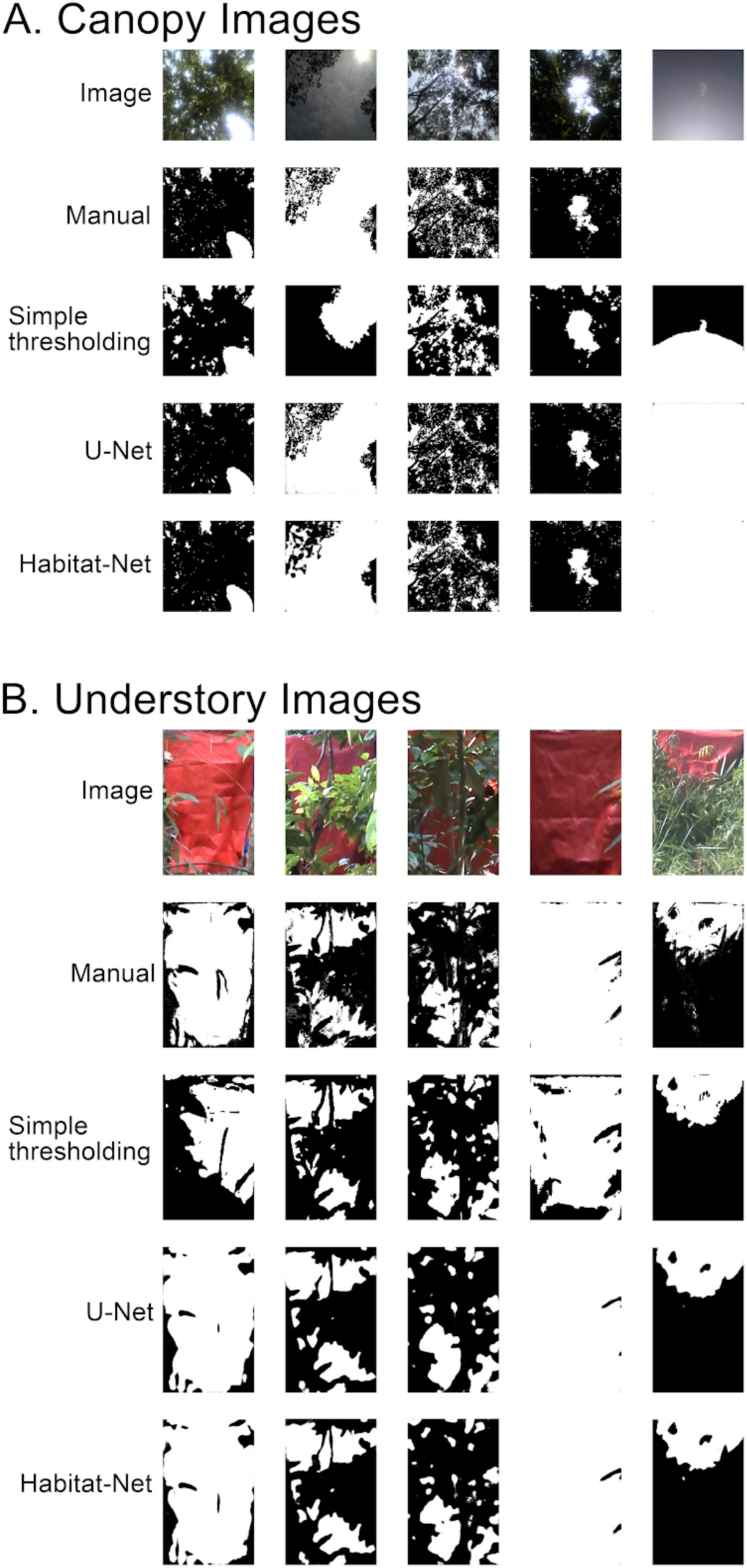
Visual comparison of the quality of segmentation rasters predicted by Simple thresholding, U-Net, and Habitat-Net to the manually segmented reference images for different situations.

**Figure 4:**
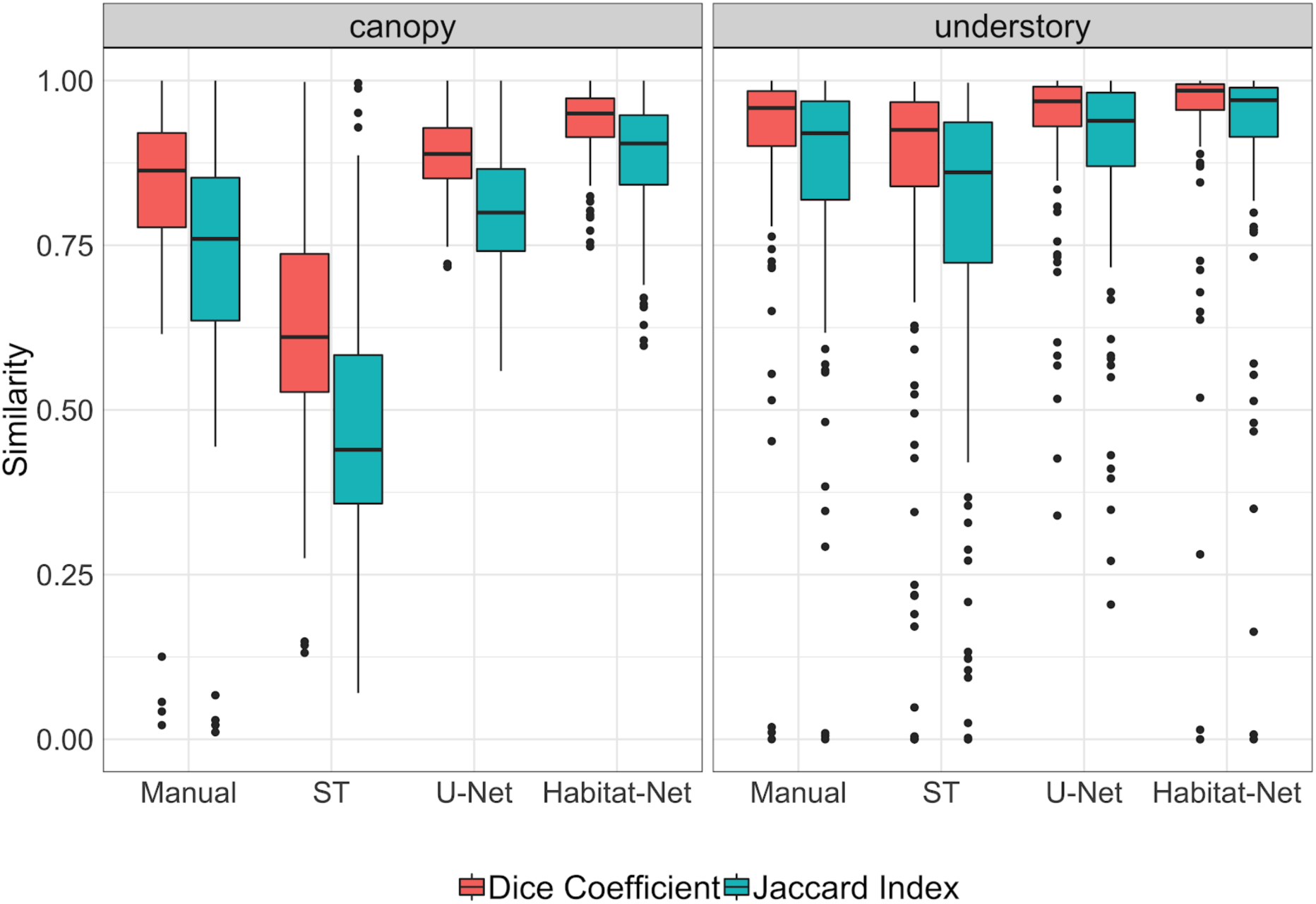
Box plots of the similarity scores (Dice coefficient and Jaccard index) between the image segmentation output by four methods and the reference manual segmentation for canopy and understory images.

## 4. DISCUSSION

Deep convolutional networks have been shown to outperform many other methods in various visual computing tasks and domains (Krizhevsky, Sutskever, & Hinton, 2012). CNNs have, however, seen little application in the ecological domain. With Habitat-Net we present an automated pipeline to process hundreds of color vegetation photographs (both canopy and understory) in a standardized, efficient and reproducible way. Our approach saves a huge amount of human labor and helps overcome the time demand associated with habitat interpretation. Habitat-Net has two advantages over manual processing, segmentation is: (1) significantly faster, and (2) more consistent. Habitat-Net produces binary segmentations with a higher similarity to the manual reference segmentation than do the approaches using simple thresholding or U-Net. For canopy images, the implementation of Habitat-Net led to a significant increase in accuracy and consistency of the image segmentation. For the understory images all methods produced a high similarity score, with Habitat-Net performing best and edging out U-Net. Habitat-Net performed well to segment images with a wide range of lighting conditions and sharpness, and the proposed method performed remarkably well even in situations where very little sky or orange flysheet was visible in a photograph (Fig. 3). The inclusion of a batch normalization layer after every convolution layer in Habitat-Net avoided internal covariate shift and enabled faster learning rates. This proved to stabilize the training (Fig. S3), boost the performance of Habitat-Net and improve the accuracy of multi-channel segmentation. Habitat-Net also outperforms previous machine learning-based methods for quantifying canopy closure. Previous research applying the ADaptive Object REcognition (ADORE; Draper, Bins, & Baek, 2000) system to canopy closure segmentation tasks produced a mean pixel-level similarity score (Dice coefficient) of 0.54 ± 0.14 (Levner & Bulitko, 2004). In contrast to canopy closure, so far there are no other automated pipelines available for the analysis of understory photographs. Therefore, Habitat-Net is the first tool that allows foresters and ecologist to describe quantitatively both the horizontal and vertical forest structures.

In this study we only quantify understory vegetation density and we did not assess vegetation complexity. However, the quantification of vegetation complexity, which provides further insight into forest structure, and thus into the ecosystem function of the understory (Halpern and Spies, 1995, Legare *et al*., 2002; Gilliam, 2007; Russell *et al.*, 2014) would also be possible using the binary raster produced by Habitat-Net. In this case, researchers would need to take many field photographs with the contrasting flysheet at different distances. As the processing of these hundreds of photographs is automated with Habitat-Net these photo series could be used to reconstruct the vegetation complexity, without the need of expensive ground-based or airborne LiDAR scans.

For ecologists, the canopy and understory habitat are important indicators of forest disturbance and, for some wildlife species, tracking these disturbances might be a warning signal of potential population declines. Furthermore, the habitat information can be combined with spatial statistics, such as species distribution models (SDMs) to assess species occurrence or abundance data (Niedballa *et al.*, 2015). This allows researchers to determine habitat associations of little known species and the knowledge gained ecological about the species can be used for more effective conservation efforts. Therefore, Habitat-Net has the potential to be an efficient and effective tool for both foresters and wildlife ecologists.

Although our network is designed with small datasets in mind, deep learning works best with large datasets (Goodfellow, Bengio, & Courville, 2016; Norouzzadeh *et al.*, 2018). Even with a relatively small training dataset available, Habitat-Net performed with a high level of accuracy. Norouzzadeh *et al.* (2018) point out that the accuracy of deep learning methods further improves as more labeled data are provided during training. Thus, as more datasets become available the performance of the network may improve by building on knowledge from multiple datasets in a process known as transfer learning (Norouzzadeh *et al.*, 2018).

A major limitation of manually processed images is the lack of standardization. The manual processing of images by humans introduces observer bias. For example, within one project a few people may do the manual processing, or in a long term monitoring program the observer (for example forestry staff) changes with time. Both of these scenarios would introduce bias and the subjectiveness inherent in the manual processing of images, making comparisons between the photographs or years difficult. Our study showed that the similarity scores between two different researchers doing the manual processing were lower than using Habitat-Net. One of the biggest strengths of Habitat-Net is, therefore, the ability to eliminate this user produced bias and standardize results which allows for inter-study site comparisons and long term monitoring.

Automated workflows, by default, accept biased inputs and can, therefore, generate undesired results. During manual processing the observer is able to sort out photos which are out of focus or of poor quality. These photos are, however, included in the automated workflow, which may lead to inaccurate estimations. Such poor quality photos could potentially be automatically removed, but this would require a high level of standardization in how the photos are taken in the field. For all photos we used the GPS unit’s built-in camera, which is a very basic camera that allows little control over camera settings. To increase standardization to a level that could potentially allow for automated removal of poor quality photos all images would have to be taken with the same camera, same settings and at the same camera angle, framing and distance. Although this would require carrying an additional higher quality camera into the field, as well as more time spent to set up equipment, we are certain that such higher quality photographs will increase the performance of Habitat-Net and, thus, the accuracy of the analysis. Currently it is not possible to automatically crop the understory vegetation photos and, thus, each image still required a minimal amount of manual processing. A satisfactory solution to do this for our images could not be found as often most of the orange flysheet was covered by vegetation making the automated recognition of the flysheet impossible. Common methods used to automatically crop images cannot be applied to our understory vegetation photographs due to the lack of common features that are always present (such as a frame). Other machine learning methods for cropping focus on “salient” image regions. The basic idea is to use information learned by the CNN about where human viewers fix their gaze to center a crop around the most interesting region (Rahman *et al.*, 2018). However, these methods are also not applicable for our understory photographs since the photographs are cluttered with no one point of interest. Such limitations could be overcome through innovative standardized practices in the field. The best solution would be to always have the flysheet centered and then perform a batch crop on all images to the specified area. This could be implemented in the field by placing a “stencil” or guide over the camera lens that is used to frame the flysheet in the center of the image. Then a technique called Object Localization and Detection could be implemented to detect the bounding box or frame (Sermanet *et al.*, 2013).

Habitat-Net provides a fast, accurate and standardized method to analyse canopy closure and understory vegetation photographs. With some optimisations during the field data collection, such as using a higher quality camera, placing a stencil over the camera lens the accuracy of Habitat-Net could even be improved and the time consuming manual cropping would not be necessary any more. With Habitat-Net, we can overcome the time lag from data collection to data processing, which often hinders timely management decisions and thus assist more sustainable forest management and conservation. Studies in other habitats might need a preliminary annotated dataset to train the model, but the overall time required to process habitat photos will be reduced and the accuracy will be increased, particularly for large projects. We hope that other users add additional datasets and their modifications of the codes to github to expand the applications and focus of Habitat-Net further. Through this a large repository of habitat images can be built, which would in turn benefit both the ecology and machine learning communities.

## Acknowledgements

We thank the Sabah Biodiversity Center and the Sabah Forestry Department, especially Johnny Kissing and Peter Lagan for support and involvement in this project. Many thanks go to the field team for their hard work to take all the habitat photographs. We would also like to thank Srijita Guha for helping with the manual cropping of understory images. This project received financial support from the German Federal Ministry of Education and Research (BMBF FKZ: 01LN1301A), Point Defiance Zoo and Aquarium through Dr. Holly Reed Conservation Fund and San Francisco Zoo.

## Author contributions

AW and AM designed the study; AV, AK, and AM wrote, trained, and test the Habitat-Net model. STW and AM collected the field data. STW, AN, AM, and SW conducted the manual image segmentation. JFA analyzed the results. JFA, AW, and AM lead writing the manuscript. All authors contributed to drafts and gave final approval for publication.

## Supplementary material

**Figure S1:**
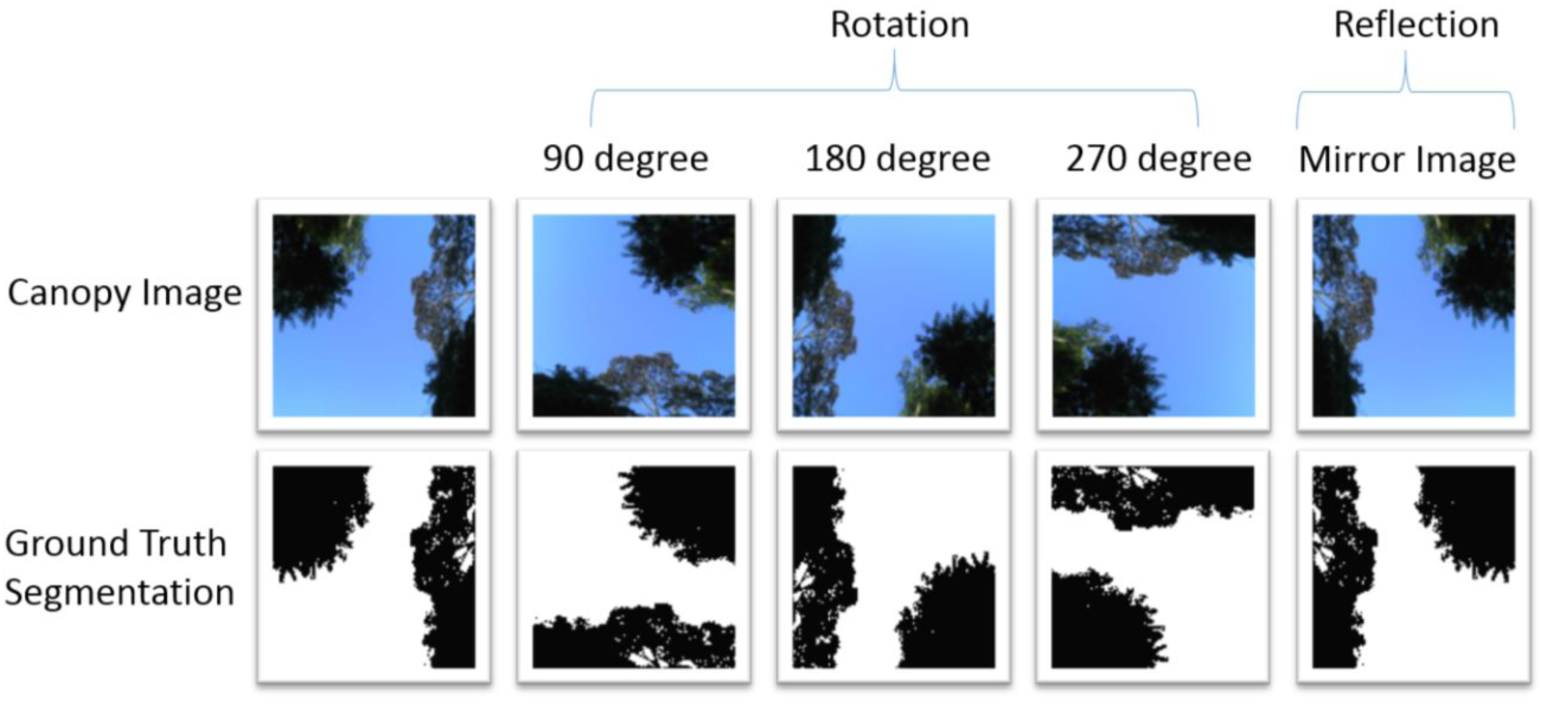
An example of data augmentation for the canopy image dataset

**Figure S2:**
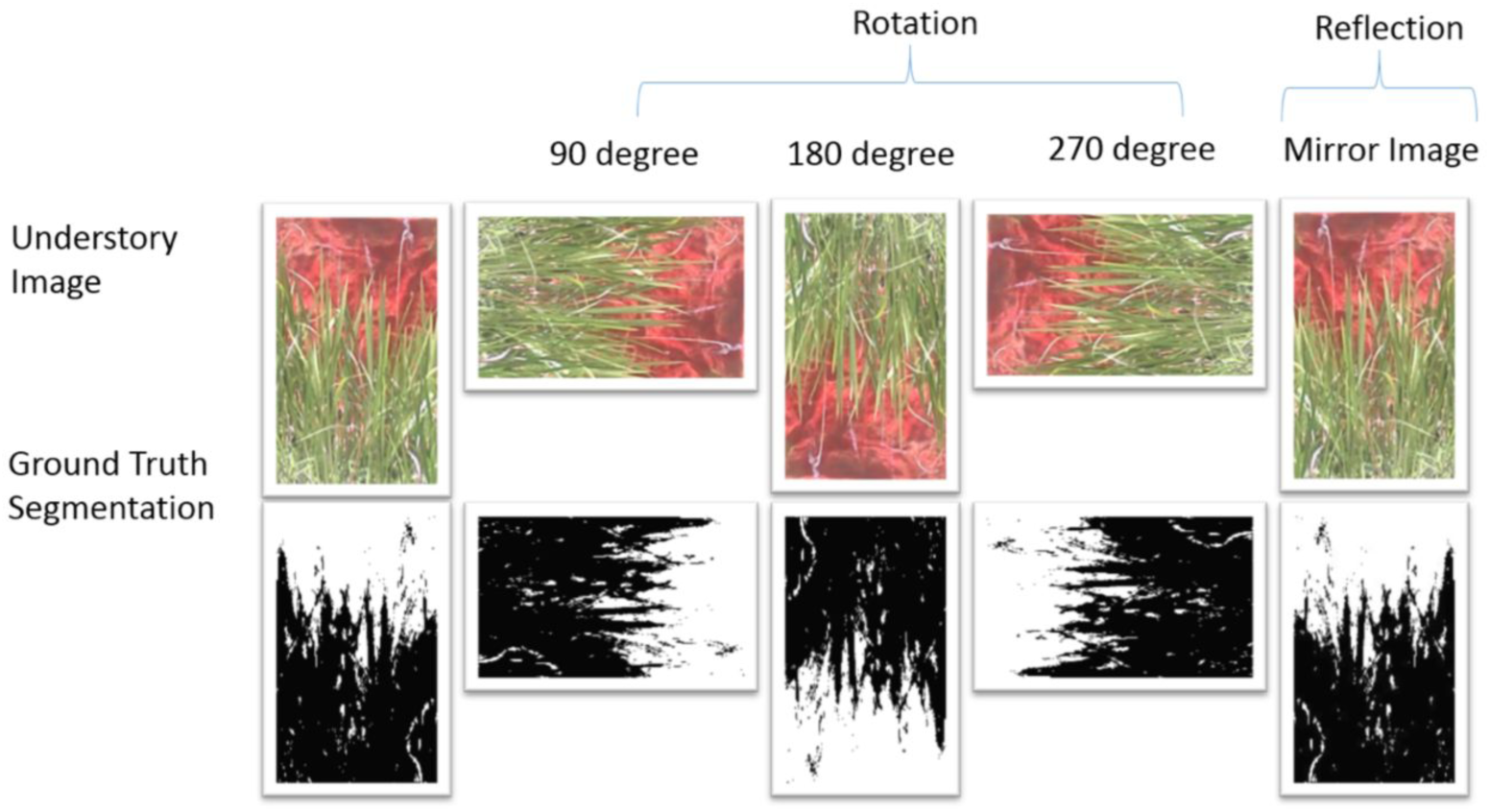
An example of data augmentation for the understory image dataset

**Figure S3:**
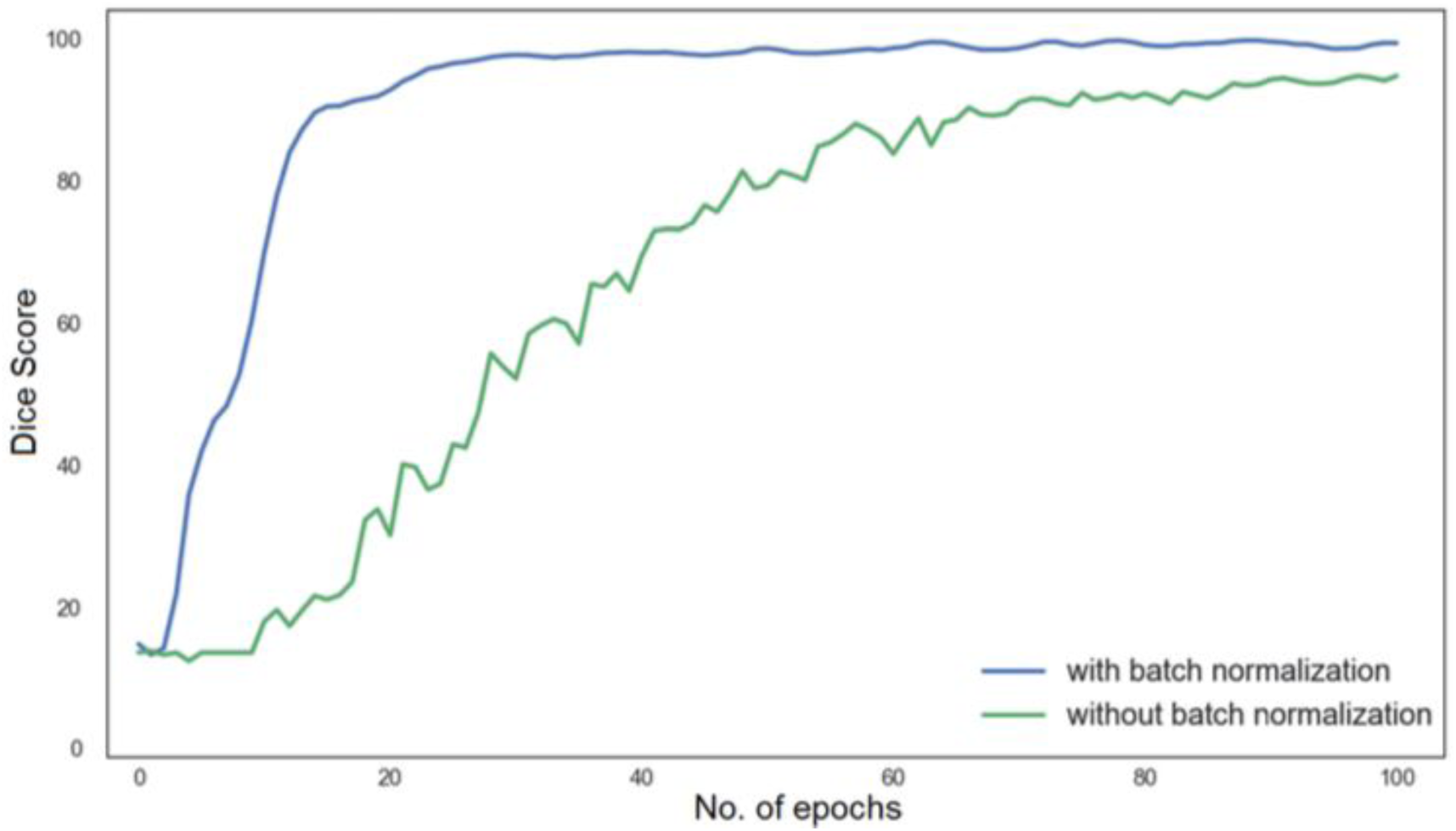
Convergence with and without batch normalization.

